# HIV-1 infection does not confer intrinsic resistance to cell death induced by cytotoxic T lymphocytes

**DOI:** 10.64898/2026.03.23.713717

**Authors:** Niklas Bachmann, Bailey Kim, Francesco R. Simonetti, Colin M. Kovacs, Rebecca Hoh, Steven G. Deeks, Janet D. Siliciano, Robert F. Siliciano

## Abstract

To eliminate the persistent reservoir of cells harboring intact HIV-1 proviruses in people living with HIV-1 (PLWH), cure strategies like the Shock-and-Kill approach rely on effector functions of cytolytic T lymphocytes (CTL). CTL are involved in the initial control of HIV-1 viremia and target productively infected cells throughout the course of infection. However, selective killing of susceptible cells could generate a reservoir dominated by cells with dysregulated cell death pathways or other features conferring resistance to killing. Here, we use CTL-engaging single-chain diabodies to assess the rate of lysis of uninfected and HIV-1-infected primary CD4^+^ T cells under identical CTL pressure in the settings of both latent and active infection. Our findings indicate that with this mode of CTL triggering, infected and uninfected CD4^+^ T cells from PLWH on ART are generally lysed at identical rates, and that an apparent survival advantage for actively infected CD4^+^ T cells primarily reflects the reduced surface antigen availability through previously described Nef-dependent downregulation of MHC class I molecules. No survival advantage is observed when the CTL response is directed through diabodies to the stably expressed non-classical MHC class I molecule HLA-E, indicating equal susceptibility to cell death.

## Introduction

Currently, an estimated 41 million people are living with Human Immunodeficiency Virus Type 1 (HIV-1) infection (1), which, if untreated, causes fatal immunodeficiency (2). Four decades of research have brought forward more than 30 antiretroviral drugs that block different steps of the viral life cycle, preventing new infection events and allowing people living with HIV-1 (PLWH) to maintain suppression of viremia to levels below the detection limit of standard clinical assays (3–5). However, a reservoir of latently infected cells persists even after many years of viral suppression (6–13). Quantification of the inducible reservoir in PLWH on antiretroviral therapy (ART) demonstrated only very slow decay, with a half-life of 44 months (10). Recent results show that after slow decay at this rate for several years, the reservoir slowly increases in size due to infected cell proliferation (12). Therefore, suppression of viral replication with ART is alone unlikely to eliminate the infected cell population, and PLWH must rely on lifelong ART to prevent viral rebound that is typically seen within weeks of treatment interruption (14–16). Additional interventions are clearly needed to target this reservoir.

The role of CD8^+^ cytotoxic T lymphocytes (CTL) in the immune response to HIV-1 is well documented (17–19): During acute infection, virus-specific CTL are instrumental in curtailing the exponential increase in viremia (20, 21). In vivo studies of Simian Immunodeficiency Virus (SIV) infection in rhesus macaques, a commonly used model for HIV-1 infection, showed that depletion of CD8^+^ T cells concurrent with virus inoculation prevented the normal post-peak reductions in viremia (22). Similarly, CD8 depletion during chronic infection led to dramatic increases in viremia (22, 23). Furthermore, the overrepresentation of certain HLA class I alleles in PLWH who maintain near undetectable viral loads without therapeutic intervention provides additional evidence for the importance of CTL in the control of HIV-1 infection (24, 25).

One widely discussed approach for curing HIV-1 infection is the Shock-and-Kill strategy targeting latently infected cells (26, 27). Latency is first reversed using pharmacologic agents so that latently infected cells can be identified by the immune system or other immunotherapeutic interventions and killed. For these approaches, it is crucial to understand the susceptibility of HIV-1-harboring cells to killing by immune effector cells. Several HIV-1 proteins affect pathways involved in cell death or survival. The HIV-1 Tat protein can sensitize human CD4^+^ T cells to apoptotic stimuli through various pathways (28–31), but also upregulate the anti-apoptotic protein Bcl-2 (32–34), which has been implicated as a relevant factor in HIV-1 persistence (reviewed in (35)). The Nef proteins of HIV-1 and SIV can inhibit proteins involved in cell death pathways (36–40) or upregulate pro-survival pathways (41, 42), but they also facilitate apoptosis in Nef-expressing cells (43, 44) and mediate FasL-dependent depletion of CTLs (45). The HIV-1 protease can promote cell death through cleavage of procaspase 8 or the anti-apoptotic BCL-2 protein (46–50). Pro-apoptotic effects have also been documented for the accessory proteins HIV-1 Vpr (51, 52) and HIV-1 Vpu (53, 54). Furthermore, CD4^+^ T cells from PLWH on ART have been reported to exhibit higher levels of Bcl-2 compared to viremic PLWH or HIV-seronegative controls (55), potentially causing higher tolerance for pro-apoptotic signaling. In addition to the direct effects of virally encoded proteins on cell death pathways, there may be preferential infection of cell populations that are less permissive for CTL-mediated apoptosis, such as those that have upregulated the anti-apoptotic genes *Bcl-XL* and *Bcl-2* (56–59) or surface molecules like OX-40 (60). The extent to which these differences allow HIV-1-infected cells to resist host cytotoxic mechanisms is unclear.

Elegant studies by Jones and colleagues have shown that after ex vivo treatment with potent latency reversing agents, subpopulations of HIV-1-infected CD4^+^ T cells carrying inducible replication-competent proviruses are not readily killed by autologous CD8^+^ T cells even though overall HIV-1 DNA levels in the target cell population were reduced (61). Measurement of total HIV-1 DNA captures both intact and defective proviruses, with the latter constituting the majority (62, 63). The authors suggested that HIV-1-specific CTL may eliminate some cells carrying defective proviruses that retain the capacity to produce viral antigens while they are unable to eliminate all cells carrying replication-competent proviruses. A subset of cells may have been selected for intrinsic resistance to killing. Consistent with this notion, they showed that the latent reservoir was predominantly located in cells expressing higher levels of the anti-apoptotic protein Bcl-2 (56). Intrinsic resistance to killing has recently been demonstrated for CD4^+^ T cell clones isolated from the reservoir (64). Interestingly, more recent studies from the same group demonstrate that HIV-1 Nef induces cytoskeletal changes in HIV-1-infected target cells that protect the cells from killing by CTL (65).

Several recent studies suggest that over long time intervals, the latent reservoir may be shaped by selective processes, leading to alterations in the integrate site landscape (66) or susceptibility to autologous neutralizing antibodies (67). The latent reservoir may arise when infected CD4^+^ T cells revert to a resting memory state that is non-permissive for viral gene expression (68, 69). It is not clear whether properties conferring resistance to CTL-mediated killing persist and remains a stable characteristic of reservoir cells. If so, selection for resistant cells could generate a reservoir with enhanced resistance to killing. To examine the extent to which the general population of HIV-1-infected cells has intrinsic resistance to CTL killing, we designed an assay to compare the susceptibility of HIV-1-infected and uninfected CD4^+^ T cells to CD8^+^ T cell-mediated lysis under otherwise identical conditions. This allowed us to determine whether infected cells have a survival advantage with respect to the CTL response. We utilized CTL-engaging single-chain diabodies with known specificities for exogenous target peptides in the context of MHC class I to measure lysis of infected and uninfected cells. We studied both freshly isolated resting CD4^+^ T cells from PLWH as well as activated CD4^+^ T cells that were acutely infected with HIV-1. Our findings indicate that, while HIV-1 Nef reduces classical MHC class I presentation on the cell surface and thereby confers some protection against HIV-1-specific CTL, most infected CD4^+^ T cells are otherwise fully susceptible to CTL-induced cytolysis triggered by diabodies.

## Results

### The rate of CTL-induced lysis of cells harboring an intact provirus is similar to the rate of lysis of uninfected cells

There are multiple potential mechanisms through which HIV-1-infected cells could exhibit lower susceptibility to CTL-mediated lysis (Figure 1). One possibility is that resistance to lysis is a stable, cell-intrinsic characteristic of a subset of CD4^+^ T cells, independent of viral infection or activation state. Reservoir cells could exhibit a resistant phenotype relative to unaffected cells if the resistance to killing was associated with other properties that allowed preferential infection (Figure 1A). Alternatively, if resistance is a stable property, then gradual selection over time could yield a population of infected cells with high resistance to lysis (61) (Figure 1B). It is also possible that resistance is a property induced by viral infection (Figure 1C). The net effect of expression of the viral proteins discussed above could be anti-apoptotic, in which case productively infected cells would be lysed at a slower rate than uninfected cells, with the difference dependent on active expression of the HIV provirus. Cells that survive and enter a state of latent infection may or may not retain this anti-apoptotic state depending on the mechanism involved.

**Figure 1.**
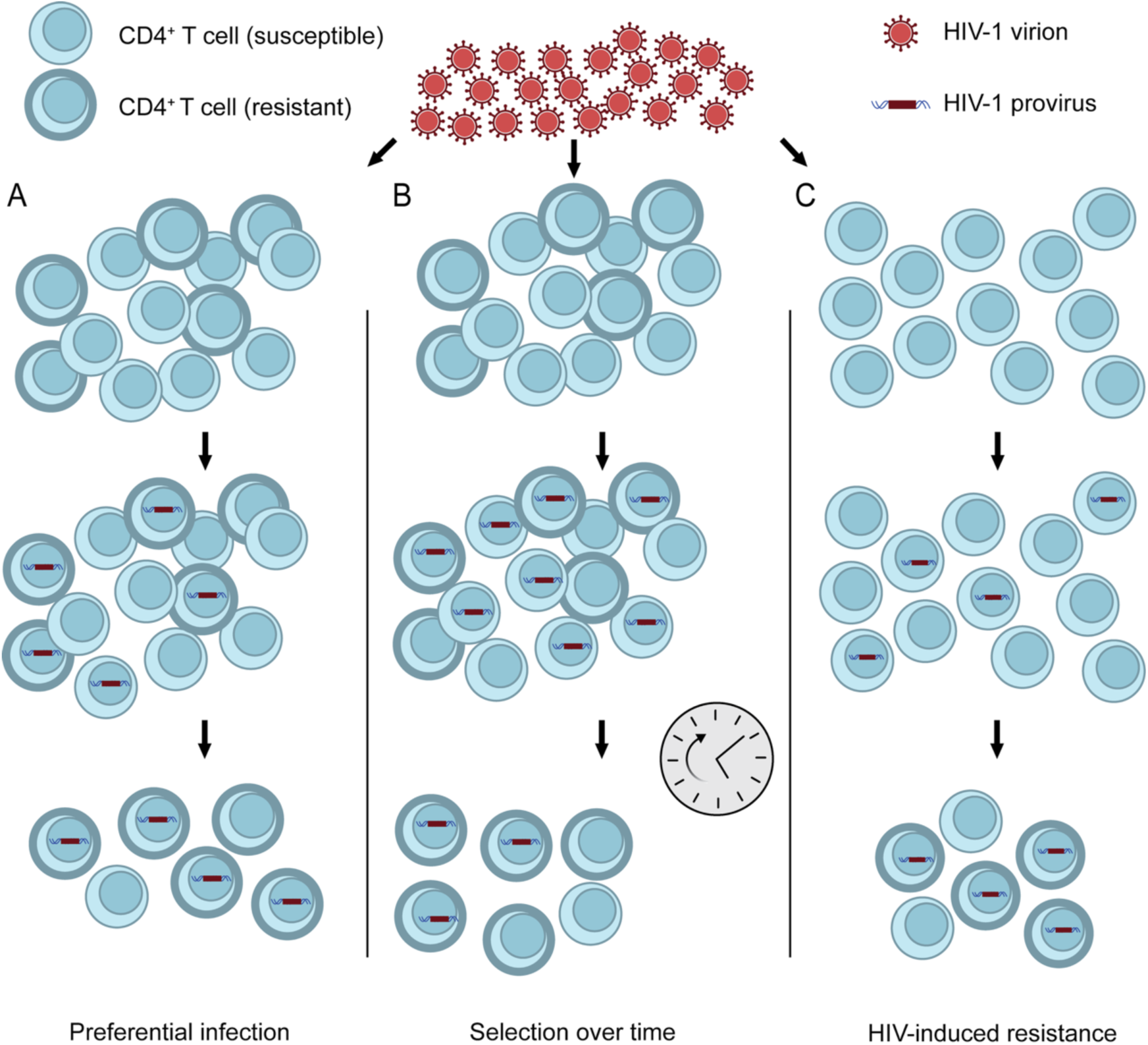
Potential mechanisms for infected cell resistance to CTL-mediated lysis. CD4^+^ T cells exhibit stable differences in susceptibility to lysis (A and B). If resistance is associated with properties that allow preferential infection, then infected cells may show greater resistance than uninfected cells (A). Alternatively, intrinsically susceptible cells may be eliminated over time, with death-resistant HIV-1-harboring cells remaining (B). A third possibility (C) is that HIV-1 induces phenotypic changes that increase cell death resistance of infected cells (HIV-induced resistance).

To address these possibilities, we designed an ex vivo assay to test for differential rates of lysis of infected and uninfected CD4^+^ T cells from PLWH on suppressive ART (Figure 2A). In short, CD4^+^ T cells were isolated from HLA-A*02:01-positive PLWH (n = 12, Table S1), and the frequencies of cells harboring an intact HIV-1 provirus were measured using the Intact Proviral DNA Assay (IPDA) (70). The CD4^+^ T cells were then pulsed with an exogenous mutant p53 peptide (p53^R175H^) that binds HLA-A*02:01, leading to presentation of the HLA-A2:p53^R175H^ complex on the cell surface (71). These target cells were then cultured together with autologous CD8^+^ T cells and a single-chain diabody that redirects CTL to target HLA-A2:p53^R175H^-presenting cells regardless of their antigen receptor specificity (Figure 2B). The next day, cells were stained for lineage, activation, and viability markers to determine the amount of CTL-mediated lysis of the total CD4^+^ T cell population (infected and uninfected cells) (Figure 2C). In this system, lysis of CD4^+^ T cells was strongly dependent on the presence of the HLA-A2:p53^R175H^- specific diabody (Figure 2D). After the co-culture, viable CD4^+^ T cells were isolated through sequential removal of dead cells and CTL based on apoptotic surface markers and lineage markers.

**Figure 2.**
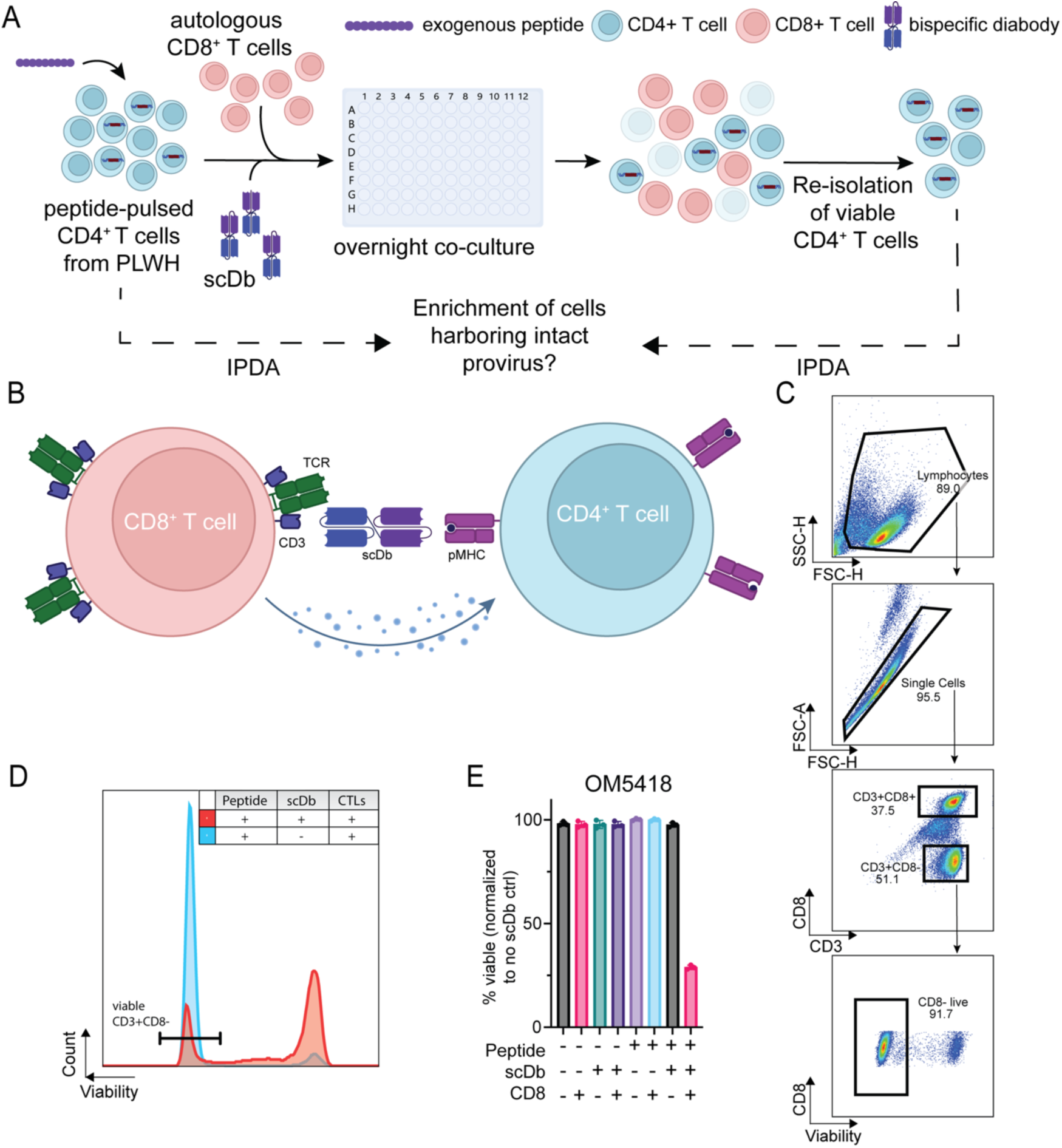
Experimental method to assess resistance to CTL-mediated killing. (A) Co-culture setup: primary CD4^+^ T cells from PLWH are pulsed with mutant p53 peptide and cultured with autologous, pre-expanded CD8^+^ T cells and p53-specific scDb. IPDA is performed on the viable CD4^+^ T cell population isolated before and after the co-culture to assess for enrichment of infected cells, indicative of resistance. (B) scDb mechanism of action: scDbs bind to cognate peptide:MHC complexes on the target cell surface and also to the CD3ε subunit of the TCR complex to induce localized cytotoxicity. (C) Flow cytometry gating strategy to determine frequency of viable target cells. (D) Representative histograms of co-cultures with and without scDbs. The reduction in the viable CD4^+^ T cell count (marked region) was used to determine the amount of CTL-mediated lysis. I Representative viability data for study participant OM5418 for the experimental condition and seven control conditions.

Genomic DNA was then isolated and assayed for intact HIV-1 proviruses using the IPDA (70). We compared the frequency of cells harboring an intact HIV-1 provirus before and after the co-culture with CTL. Control co-cultures demonstrated that killing of CD4^+^ T cells in this system was dependent on the presence of the p53^R175H^ peptide, the diabody, and CD8^+^ CTL (Figure 2E).

In the coculture system described above, CD8^+^ CTL are redirected to kill targets cells based on target cell expression of HLA-A2:p53^R175H^ complexes, not HIV-1 antigens. Thus, both infected and uninfected cells should be recognized and lysed to the same extent if latently infected cells have no general intrinsic resistance to CTL killing. If infected cells are lysed at a lower rate than uninfected cells due to intrinsic resistance, enrichment of intact proviruses in the surviving population is expected. We defined enrichment (E) as the log_10_ of the ratio of final over initial IPDA frequencies:

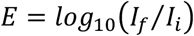

where 𝐼_𝑓_ and 𝐼_𝑖_ are the final and initial IPDA values. Intrinsic resistance should be evident as 𝐸 > 0. It is also possible that infected cells may be more susceptible to lysis than uninfected cells, which would have the opposite effect, decreased IPDA values after the coculture (𝐸 < 0).

Cells from each study participant were tested in at least two independent assays. The frequencies of intact proviruses before and after culture were in the range observed previously in a large-scale study of PLWH on ART (72). Contrary to the notion that infected cells have intrinsic resistance to killing, we found that in 22 out of 25 assays, the frequency of cells harboring an intact provirus remained constant or decreased throughout the co-culture, indicating no significant enrichment (p-value 0.3126, paired two-tailed t-test; Figure 3B), despite reduction of the target cell population of up to 63% percent (average across all individual assays 38.0 ± 22.9%). Only 3 of 25 assays showed an enrichment of >0.5 logs, although these increases were not consistently observed in multiple assays from the relevant participants. For donor SCOPE1597, two assays showed a substantial increase, while a third one showed a slight decrease in the frequency of intact proviruses. On average for the 3 assays, we detected 3.10 and 21.0 copies per million CD4^+^ T cells before and after the co-cultures for this donor, respectively, with a mean lysis rate of approximately 13.5%. It is worth nothing that the baseline frequency of intact proviruses in this donor was near the limit of detection based on intact proviral frequency and the number of cells available for analysis, and in two out of six measurements, one before and one after a co-culture, we did not detect any intact provirus in 1.2 x 10^6^ cells and 0.51 x 10^6^ cells, respectively. Similarly, in the case of participant SCOPE2243, one assay showed no detectable intact provirus in 1.2 x 10^6^ cells before or 1.1 x 10^6^ cells tested after the culture, whereas we observed an increase from 15.1 to 228.3 copies per million cells in another assay (average across assays: 7.95 vs 114.6 intact copies per million cells), with a mean of 62.8% target cell lysis.

**Figure 3.**
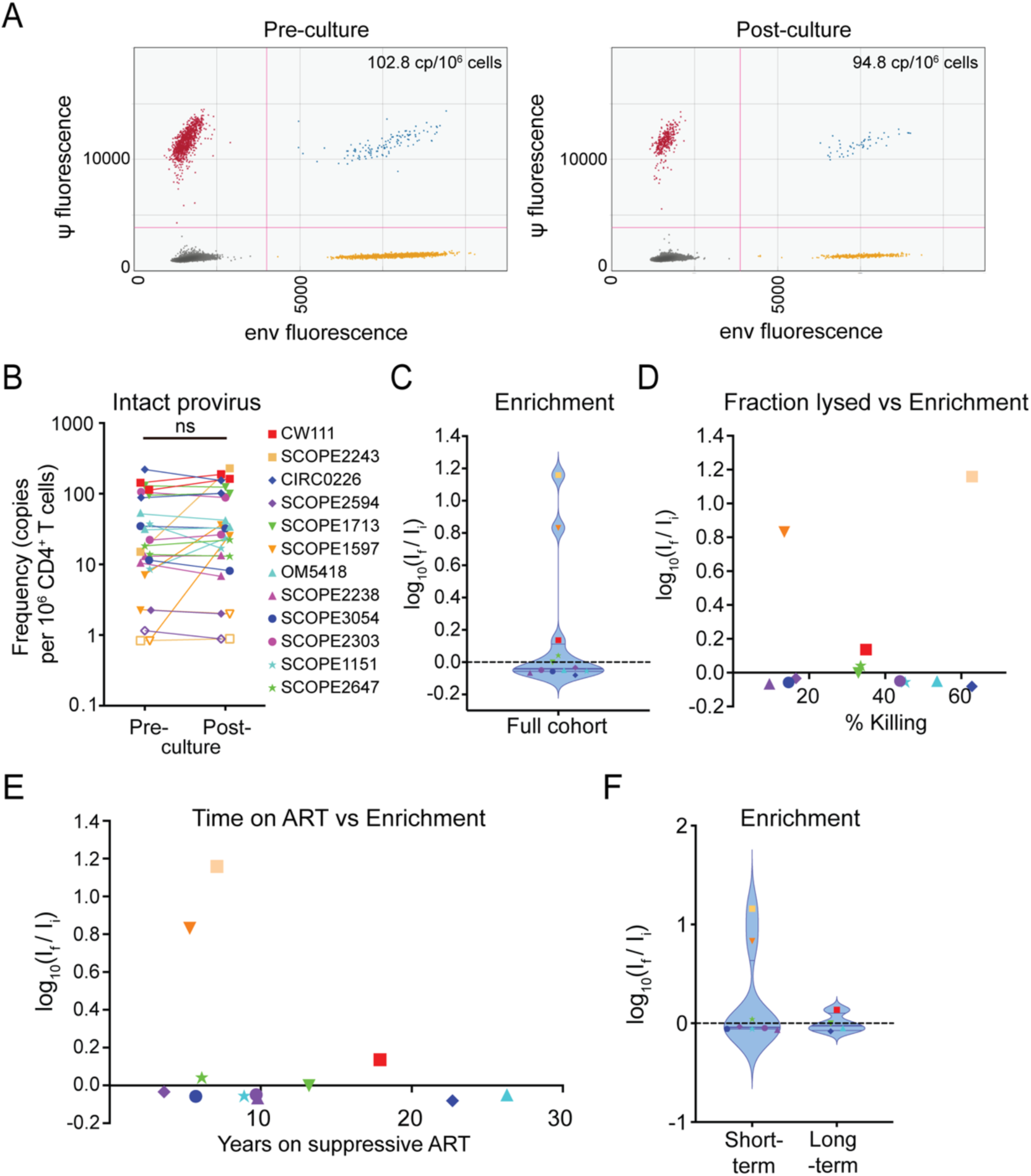
Ex vivo co-cultures with cells from PLWH do not show enrichment for infected cells in ten out of twelve participants. (A) Representative ddPCR plots of IPDA performed on CD4^+^ T cells from donor CIRC0226 before and after co-culture. Each dot represents one droplet out of at least 10,000 droplets measured per assay. Measured frequencies were corrected for cell count and DNA shearing as described previously (70). (B) Pre- and post-culture frequency of intact provirus per 1x10^6^ CD4^+^ T cells. Each point represents the calculated frequency of intact provirus for one assay for the indicated study participant. (C) Violin plot of enrichment for total study cohort. Each dot represents the average enrichment score for an individual study participant. Colors and symbols coding as in (B). Dashed line marks no change in frequency; horizontal lines mark the median and first and third quartile. (D) Correlation between extent of killing and enrichment for intact proviruses. (E) Correlation between time on ART and enrichment of intact provirus. (F) Comparison of enrichment of intact provirus between study participants with less than 10 years of continuous ART (short-term) and over 13 years of continuous ART (long-term).

Figure 3C shows a plot of enrichment values for 12 participants averaged over at least two independent experiments. The mean value for E was not significantly different than zero (p= 0.23, two-sided t-test compared to a theoretical mean of 0), indicating no general enrichment of intact proviruses. In principle, enrichment is easier to observe when a higher fraction of the CD4^+^ T cells is lysed. We therefore examined whether the observed degree of enrichment was related to the fraction of CD4^+^ T cells lysed in the coculture (Figure 3C). Notably, enrichment values were not clearly correlated with the degree of killing (Pearson correlation coefficient r = 0.15, p = 0.63).

One hypothesis for intrinsic resistance to CTL-mediated lysis in HIV-1-harboring cells is the selective pressure that leads to the enrichment of resistant cells over long time intervals (Figure 1B). However, we did not observe a positive correlation between the time on ART and the relative enrichment of cells harboring intact provirus in our assay (Figure 3E, Pearson Correlation coefficient r = -0.31, p = 0.33). Furthermore, Figure 3F shows a direct comparison of study participants on short-term ART (< 10 years, average 7.0 years, n = 8) and long-term ART (> 13 years, average 20.0 years, n = 4). We did not observe a substantial increase in enrichment throughout the co-culture in the long-term cohort (average frequency of intact provirus 109.3 vs 113.2 copies/10^6^ CD4^+^ T cells, E = 0.015) compared to the short-term cohort (18.81 vs 32.80 copies/10^6^ CD4^+^ T cells, E = 0.244).

In summary, infected CD4^+^ T cells from PWH on ART do not generally show measurable intrinsic resistance to CTL killing.

### CD3-engaging scDbs exhibit some latency reversing potential

In PLWH on ART, most proviruses persisting in circulating CD4^+^ T cells are in a latent state of infection reflecting the resting state of the host cells (73). In experiments described above, the cells were not treated with any activating stimuli or latency reversing agents, and no general resistance to cytolysis was observed. However, it remains possible that a state of resistance is dependent on active viral gene expression. We therefore considered the possibility that the CD3-binding domain of the scDb might induce some degree of activation of CD4^+^ T cells in the cultures. As CD4^+^ T cells also express the same CD3 surface subunit that is engaged by the scDbs on CTLs, we assessed scDb-dependent upregulation of the early activation marker CD69 on CD4^+^ T cells at the end of the co-cultures described above. Surface staining and flow cytometric analysis confirmed an increase in the frequency of CD69^+^ CD4^+^ T cells (Figure 4A: 4.2% vs 20.9% of CD4^+^ T cells, p-value < 0.000001). To test whether the scDb-mediated activation of primary CD4^+^ T cells from PLWH leads to increased levels of viral transcription, we performed a digital PCR (dPCR)-based viral quantitation assay (VQA) (Figure 4B) using total CD4^+^ T cells isolated from three study participants with larger HIV-1 reservoirs (average 100.4 ± 52.9 intact proviruses/10^6^ CD4^+^ T cells). Cells were pulsed with either an irrelevant peptide (HLA class I signal peptide, VMAPRTLVL) or the p53^R175H^ mutant peptide following the same protocol as for the cytotoxicity assays, and then cultured for 18 hours in the presence of the HLA-A2:p53^R175H^-specific scDb at a concentration of 5 nM. As a positive control, cells were treated with phorbol 12-myristate 13-acetate (PMA, 50 ng/ml) and ionomycin (1 μM), a combination that induces strong T cell activation (74). In negative control wells, cells were cultured with media only. Cell-associated RNA was then isolated and reverse transcribed using oligo-dT and random hexamer primers, and the levels of different HIV-1 transcript species were measured by dPCR. We used labeled probes complementary to a sequence in the integrase region of the HIV-1 *pol* gene (int), to the splice junction for the HIV-1 *vpu-env* spliced transcript, or to the terminal nucleotides and poly-A tail to assess levels of unspliced, spliced, and polyadenylated HIV-1 RNA (75). Measured transcript levels were normalized to the amount of input RNA, as measured by Qubit. Each tested culture condition had three replicate wells, which were then each tested in two technical replicates for the dPCR reaction.

**Figure 4:**
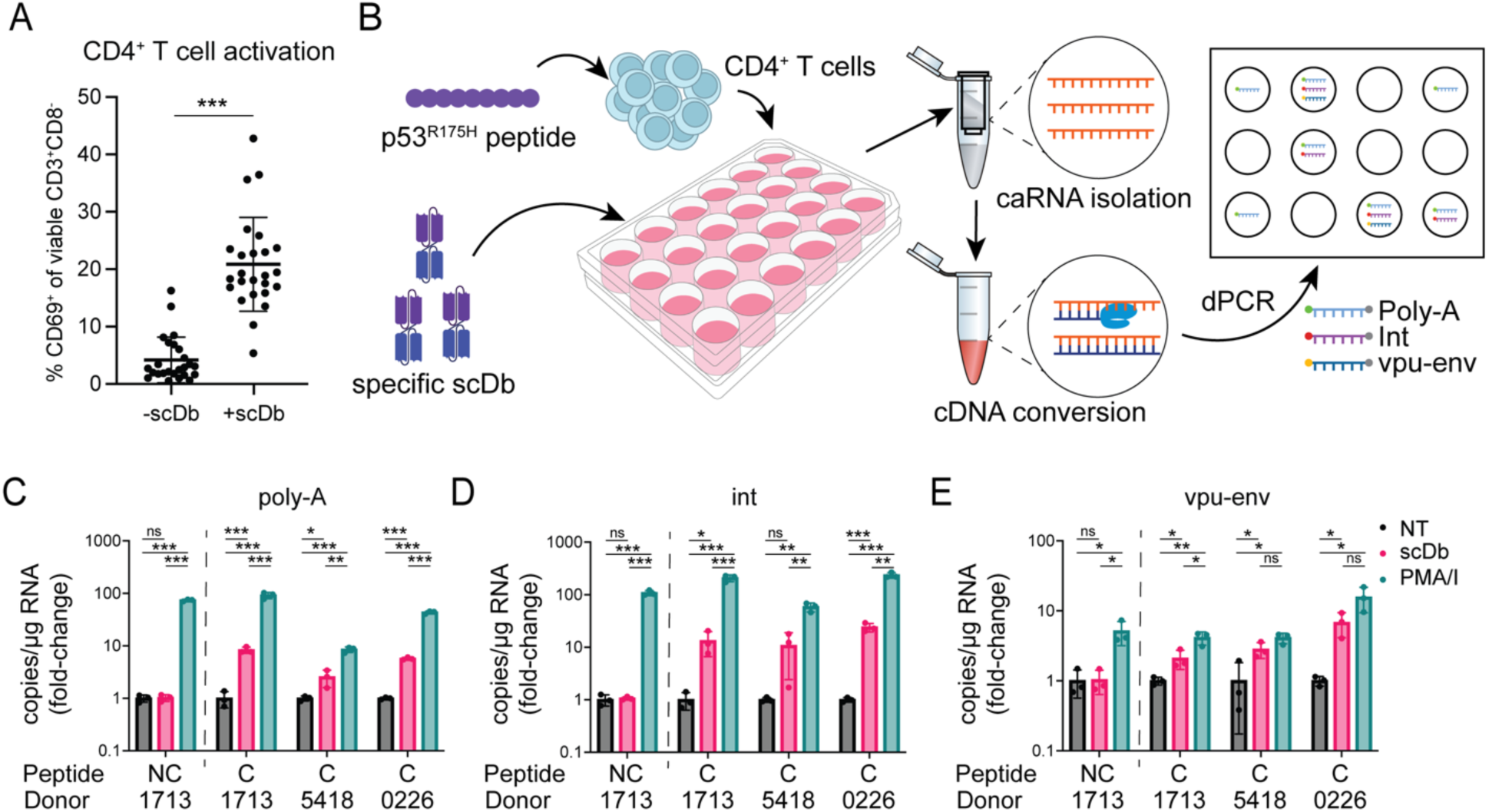
pMHC-specific latency reversal by the p53-specific scDb. (A) Frequencies of CD69^+^ CD4^+^ T cells in individual co-cultures without and with scDb across at least two co-cultures for all twelve study participants. Co-cultures were set up as described in Figure 2. Error bars represent standard deviation. (B) Induction of viral RNA by the p53-specific scDb. Primary CD4^+^ T cells from HIV-1-positive study participants were pulsed with the p53^R175H^ peptide (n = 3) or an irrelevant control peptide (HLA-A2 signal peptide, VMAPRTLVL; n = 1) and cultured for 18 hours with or without p53-specific scDb. RNA was isolated and reverse transcribed. Digital PCR was performed with probes binding to poly-adenylated HIV-1 RNA (poly-A), the integrase region of HIV-1 *pol* (int) or the splice junction for *vpu-env* mRNA. (C-E) pMHC-specific increase in HIV-1 transcripts induced by the p53-specific scDb. Measurements were averaged over two technical replicates for each of three identical culture wells per condition, and normalized to transcript level in the no treatment (NT) control. PMA/ionomycin was included as positive control. Colors represent treatment conditions: black: no treatment; red: p53-specific scDb; green: PMA/I. NC: non-cognate peptide (HLA-A2 signal peptide SP-2A, VMAPRTLVL), C: cognate peptide (p53^R175H^, HMTEVVRHC).

As expected, all three assays showed a significant increase in cell-associated HIV-1 RNA following treatment of CD4^+^ T cells with PMA/I, regardless of which peptide the cells were pulsed with (fold-change for the *gag* probe = 58.37 – 235.47, for the *vpu-env* probe = 4.10 – 15.59, for the poly-A probe = 8.49 – 92.00) (Figure 4, C-E). In the presence of the p53-specific scDb, cells pulsed with irrelevant control peptide exhibited no change in transcript levels for any of the three probes (fold-change for *gag* probe = 1.061, for *vpu-env* probe = 1.032, for poly-A probe = 1.023). However, cells pulsed with the cognate p53^R175H^ peptide showed higher levels of each transcript species for all three study participants compared to untreated cells (fold-change for the *gag* probe = 10.65 – 23.98, for the *vpu-env* probe = 2.09 – 6.78, for the poly-A probe = 2.52 - 8.34), though not to the same extent as cells treated with PMA/I. This observation indicates that CD3-engaging scDbs can induce viral transcription in latently HIV-1-infected cells harboring inducible provirus in a pMHC-specific manner. In addition to direct engagement of HIV-1-harboring cells, cytokines released by activated bystander cells may further act as latency reversing stimulus. Nevertheless, no consistent difference in the susceptibility to CTL-mediated lysis was observed in our ex vivo assays.

### Induction of resistance by viral gene expression

Although the above experiments did not provide evidence for a general state of resistance to CTL-mediated killing in CD4^+^ T cells from PLWH on ART, it remained possible that resistance is dependent on high levels of viral gene expression, higher than those induced by the scDb. Ex vivo analysis of the problem is complicated by the very low frequency of CD4^+^ T cells harboring an intact provirus (typically ranging from 1 in 1x10^4^ to 1 in 1x10^6^ cells) (76), the toxicity of most latency reversal agents (77) and the fact that many latent proviruses are not induced even with global T cell activation (78–80). Therefore, to test susceptibility to CTL-mediated lysis in cells that are actively expressing HIV-1 proteins, we conducted a modified assay with acutely infected CD4^+^ T cells from five HIV-1-seronegative donors (Figure 5A). One potential mechanism by which productively infected cells can resist CTL killing is through Nef-mediated downregulation of classical MHC class I molecules (81, 82). To account for this possibility, PHA-activated CD4^+^ T cells were infected with one of three eGFP-expressing HIV-1 reporter viruses (Figure 5B).

**Figure 5:**
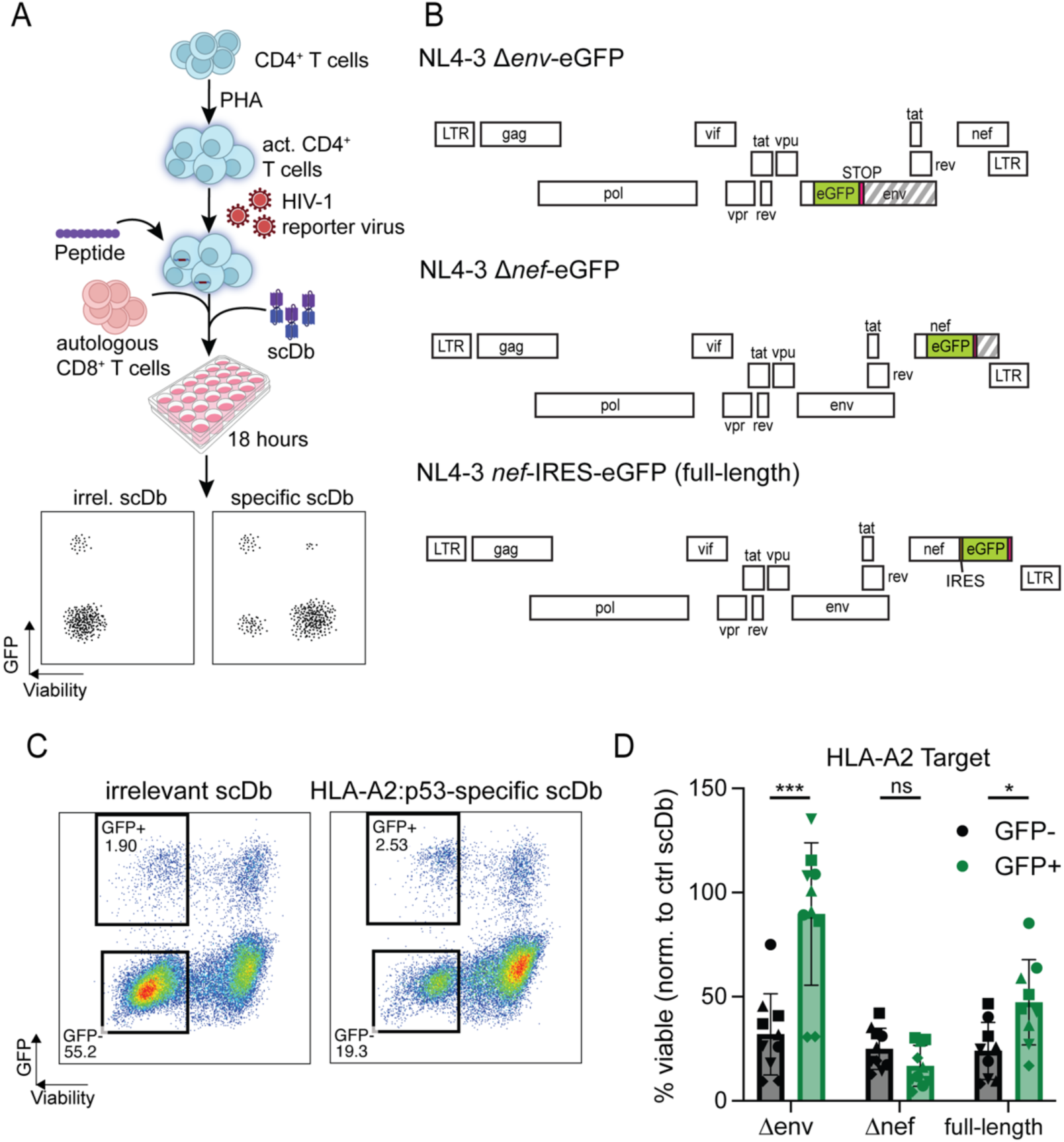
Effect of active viral expression on cell death resistance. (A) Experimental setup: Healthy donor CD4^+^ T cells were activated for 72 hours with PHA and infected with different HIV-1 GFP-reporter constructs. Cells were then pulsed with exogenous p53^R175H^ peptide and co-cultured with pre-expanded autologous CD8^+^ T cells and p53-specific scDb or irrelevant control scDb. Viability and GFP-expression were analyzed through flow cytometry. (B) Genome maps of the HIV-1 reporter constructs used to infect target cells. Each row represents one reading frame. Striped segments indicate gene segments that are no longer translated due to insertion of a STOP-codon. (C) Representative flow plot showing viable uninfected (GFP-negative) and HIV-1-expressing (GFP-positive) cell populations. (D) Viability of uninfected target cells or cells infected with indicated HIV-1 reporter constructs after co-culture with the HLA-A2:p53^R175H^-restricted scDb. Each dot shows the average of three technical replicates per condition. Symbols represent individual donors; assay was performed 2 independent times for each of the n=5 healthy donors, except for the full-length condition, where one donor was only tested once for technical reasons. A p-value less than 0.05 was considered significant. Abbreviations: PHA, phytohemagglutinin; LTR, long terminal repeat; IRES, internal ribosome entry site. ns: no significance (p > 0.05).

NL4-3 Δ*env-*eGFP (Δenv) contains an *env* gene truncated by an inserted eGFP-cassette, but encodes for all other viral proteins, and was pseudotyped with dual-tropic HIV-1 envelope (strain 89.6). In NL4-3 Δ*nef-*eGFP (Δ*nef*), the *nef* gene is truncated by the inserted eGFP-cassette.

NL4-3 *nef*^_^IRES-eGFP is a replication-competent provirus with an IRES-eGFP inserted downstream of *nef*. Three days after infection, cells were pulsed with the p53^R175H^ peptide and plated for co-culture with autologous CD8^+^ T cells in the presence of either the p53-specific scDb or an irrelevant control scDb RLP13, which recognizes the *M. tb*-derived peptide RLPAKAPLL (*Mtb44*) in the context of HLA-E, but contains an identical CD3ε-binding domain for CTL activation. After 18 hours of co-culture, the viability of cells expressing the HIV-1 constructs, identified by means of the eGFP-reporter, was compared to the viability of the GFP-negative populations across the different conditions (Figure 5C).

A representative experiment is shown in Figure 5C. Following infection with the NL4-3 Δ*env-*eGFP pseudovirus, GFP^-^ target cells are readily lysed in a p53-specific manner while productively infected cells (GFP^+^) in the same cultures were protected. However, the observed protection was *nef-*dependent. In co-cultures with CD4^+^ T cells that were infected with the Nef-expressing NL4-3 Δ*env*-eGFP and NL4-3 *nef*-IRES-eGFP constructs, there was significant enrichment for productively infected, GFP^+^ cells while GFP-negative cells in the same cultures were more susceptible to killing. In contrast, when target cells were infected with the NL4-3 Δ*nef*-eGFP constructs, GFP^+^ and GFP^-^ cells were lysed at similar rates (Figure 5D). Together, these results show that HIV-1 *nef*-expressing CD4^+^ T cells are lysed at significantly lower rates than uninfected cells or cells expressing *nef*-deficient provirus when exposed to an HLA-A2-directed CTL response.

### HLA-A2 is downregulated on the surface of actively infected cells in vitro

The apparent HIV-1 Nef-dependent effect on CTL-mediated lysis of target cells could reflect a net anti-apoptotic phenotype induced by Nef (37–41), allowing cells to tolerate exposure to cytolytic effector mechanisms without cell death. Another possibility is the previously documented (81, 82) Nef-mediated downregulation of surface HLA-A and HLA-B molecules (reviewed in (83)). Downregulation of surface HLA-A2 molecules presenting the cognate p53^R175H^ peptide for recognition by the p53-specific scDb could explain the observed protection of cells infected with *nef*^+^ viruses. To study the effect of HIV-1-infection on surface levels of HLA-A2, we infected PHA-activated CD4^+^ T cells from HIV-1-seronegative donors with HIV NL4-3 Δ*env*-eGFP or NL4-3 Δ*nef*-eGFP and then after three days sorted the cells into GFP-negative and GFP-positive fractions. The cells were then stained with lineage and viability markers as well as phycoerythrin (PE) conjugated anti-HLA-A2 monoclonal antibodies for analysis by flow cytometry. The mean fluorescence intensity (MFI) of the PE signal was used as a readout of surface levels of HLA-A2. Unstimulated CD4^+^ T cells were included for comparison. Across all tested healthy donors (n = 3), we observed an average 2.97-fold (range 1.99 – 3.87) upregulation of surface HLA-A2 in the activated, GFP-negative fraction compared to baseline level measured in unstimulated cells. However, for the PHA-activated cells, the MFI in the GFP-positive population was reduced dramatically relative to uninfected (GFP-negative) cells in the same culture and was on average 40% lower than the level on unstimulated cells (Figure 6A). In the population infected with the NL4-3 Δ*nef*-eGFP construct, surface HLA-A2 levels on the GFP-positive population did not differ from those on the GFP-negative population. This Nef-dependent downregulation of surface HLA-A2 could explain the observed survival advantage of cells infected with HIV-1 *nef*-encoding reporter viruses. These findings are in accordance with a recently published study showing protection of HIV-1 *nef*-expressing CAR T cells from alloreactive host CTL responses, which was linked to a moderate downregulation of surface HLA-A and HLA-B on *nef*-expressing cells (84).

**Figure 6:**
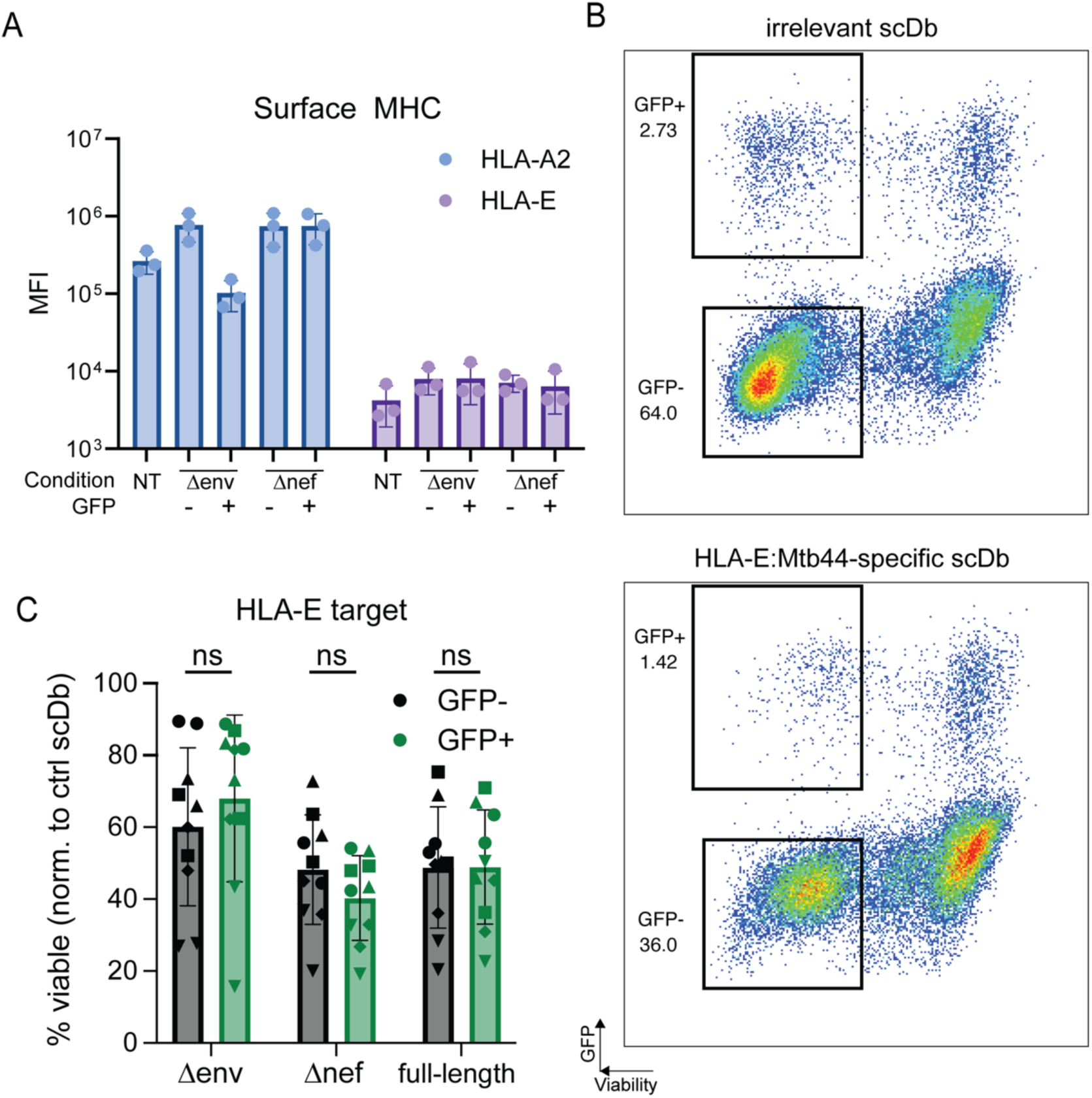
Modified assay targeting stably expressed HLA-E. (A) Cell surface levels of classical HLA-A2 and non-classical HLA-E molecules on untreated, uninfected and HIV-1-expressing cells. Healthy donor CD4^+^ T cells were either left untreated (NT) or activated with PHA over 72 hours and infected with indicated reporter virus construct. Cells were then sorted by GFP-expression and stained with PE-labeled monoclonal antibodies for HLA-A2 or HLA-E. Each bar represents average measurements across n=3 healthy donors. (B) Viability of uninfected target cells or cells infected with indicated HIV-1 reporter constructs after co-culture with the HLA-E-restricted RLP-13 scDb. Each dot shows the average of three technical replicates per condition. Symbols represent individual donors; assay was performed 2 independent times for each of the n=5 healthy donors. Bars indicate average viability and standard deviation.

### No differential susceptibility to CTL-mediated lysis mediated by scDbs targeting HLA-E

While lower antigen availability could explain the observed enrichment for HIV-1 *nef*-expressing cells, it is possible that Nef confers additional pro-survival characteristics, such as the upregulation of *bcl-2* or alterations in the cytoskeleton (65). Therefore, we repeated the surface HLA measurements described above using a PE-conjugated monoclonal antibody to the non-classical MHC class I molecule HLA-E (Figure 6A). While surface levels of HLA-E in activated cells increased slightly relative to resting cells (MFI 7951.3 vs 4206.7), there was no significant difference between HLA-E levels on the GFP-positive and GFP-negative cells for both the NL4-3 Δ*env*-eGFP and the NL4-3 Δ*nef*-eGFP conditions (Figure 6A). Thus, targeting cells for lysis using an scDb specific for a peptide presented on HLA-E would allow for comparison of susceptibility to CTL-mediated lysis without the complication of reduced HLA-A2 expression. We therefore set up a modified killing assay following a similar protocol to the one described in Figure 4A but using an *M. tb*-derived peptide *Mtb44* which binds HLA-E with high affinity (85) and is recognized by the *Mtb44*-specific scDb RLP-13. When targeting this HLA-E pMHC complex, we did not observe statistically significant enrichment for provirus-expressing cells infected with any of the of the three HIV-1 reporter construct across 5 healthy donors (Figure 6, B and C). Therefore, we conclude that the survival advantage seen when targeting HLA-A2 can be explained by the lower surface levels of classical MHC class I such as HLA-A2 and that actively infected cells generally do not otherwise show lowered susceptibility to CTL-induced cell death.

## Discussion

The main obstacle to an HIV-1 cure is the stable, long-lived reservoir of CD4^+^ T cells harboring intact HIV-1 proviruses that persists in PLWH even after years of suppressive antiretroviral therapy and that can give rise to viral rebound within weeks of treatment interruption (4–13). The role of CD8^+^ CTL in the control of both acute and chronic infection has been thoroughly documented (reviewed in (17, 86)), as has their potential to contribute to viral suppression in PLWH who control viral replication the absence of ART (25). This potential is critical to the Shock-and-Kill approach for HIV-1 cure, which generally involves pharmacologic induction of viral expression in latently infected cells and subsequent lysis by cytotoxic effector cells (26).

However, this strategy requires HIV-1-infected cells to be susceptible to cell lysis. Given the long half-life of the latent reservoir (10–12), it is possible that HIV-1-harboring cells exhibit some level of intrinsic resistance to cell death, either inherently or induced by HIV-1 infection. As discussed above, several HIV-1 proteins have been reported to dysregulate cell survival and cell death pathways, in some cases causing cytotoxicity or increasing sensitivity to apoptotic stimuli and in other cases promoting pro-survival phenotypes. Here, we assess the susceptibility of latently infected and productively infected primary CD4^+^ T cells to killing by CD8^+^ T cells and compare the susceptibility to that of uninfected cells present in the same cultures. In ten out of twelve study participants on ART, freshly isolated CD4^+^ T cells harboring intact HIV-1 provirus were lysed at the same rate as uninfected cells, indicating that generally there is no significant general intrinsic resistance to CTL-mediated killing in HIV-1-infected cells. We further show that in in vitro infections of CD4^+^ T lymphoblasts, there is Nef-dependent reduction of surface HLA-A2 complexes which correlates with a significantly reduced rate of CTL-mediated cell death in this population compared to uninfected cells. However, this effect was not observed when the killing was redirected with scDbs targeting HLA-E complexes, which are not downregulated by Nef, indicating that HIV-1-expressing cells are not inherently more resistant to CTL-mediated lysis. Rather they can partially evade recognition by limiting surface antigen availability. These findings help elucidate the net effect of HIV-1 gene products on the susceptibility to CTL-mediated killing and can inform new strategies for the immunotherapeutic targeting of HIV-1-infected cells.

It is important to note that cell death, and apoptosis in particular, is a complex process that relies on a balance of several factors rather than a binary switch. While our observations indicate that relative to uninfected cells, HIV-1-infected CD4^+^ T cells do not show a general increase in resistance to CTL-mediated killing, several studies have shown dysregulation of cell death pathways. Specifically the anti-apoptotic Bcl-2 protein was shown to be downregulated (87) or upregulated in subpopulations of HIV-1-harboring cells (56). More recent studies show that HIV-1-harboring cells were sensitized to the Bcl-2 antagonist ABT-199 (venetoclax) (56, 88), which is being further investigated in a Phase I/II clinical trial (ClinicalTrials.gov ID NCT05668026).

These observations are in line with numerous studies indicating both pro- and anti-apoptotic effects of HIV-1 infection: Upregulated Bcl-2 could act as a “brake” to prevent cell death caused by pro-apoptotic effects of HIV-1-infection. Removing this brake tips the balance towards a pro-apoptotic state, leading to ABT-199-induced cell death even in the absence of additional pro-apoptotic stimuli. However, our findings show that this dysregulation is not sufficient to withstand exposure to CTL responses ex vivo.

This study aimed to assess whether HIV-1 infection is generally associated with resistance to cell lysis through cytotoxic effector cells. While our results indicate that infected cells are overall lysed at the same rate as uninfected cells unless surface antigen availability is limited, it is likely that individual CD4^+^ T cell clones differ with respect to susceptibility. These differences could reflect integration site effects that further alter expression patterns of genes involved in antigen presentation or cell death pathways, or through mutations altering target epitopes. Additionally, a recent study from Leyre and colleagues (65) proposes an HIV-1 Nef-dependent reduction in cortical stiffness of CD4^+^ T cells through perturbation of cytoskeletal dynamics, on top of pre-existing heterogeneity in this population. Since signal transduction through the TCR has previously been shown to depend on mechanical forces (89, 90), some infected cells could evade CTL killing due to insufficient stiffness for mechanosensing. Interestingly, this protective effect is bypassed by the CD3-engaging diabodies used in this study. Therefore, it is possible that a subpopulation of cells harboring HIV-1 proviruses may withstand exposure to cytotoxic host responses simply because they fail to induce TCR-dependent degranulation. Additional experiments assessing CTL degranulation in response to these subpopulations could help elucidate their mechanisms of survival.

Our study has limitations. The study cohort included 12 individuals, with eight participants classified as short-term treated (3.6 – 9.8 years on suppressive ART) and four participants classified as long-term treated (13.2 – 26.3 years on suppressive ART). While 11 of the 12 study participants showed no consistent and substantial enrichment for cells harboring intact provirus in multiple independent assays, participant CW111 showed a slight but consistent enrichment of approximately 0.14 logs or 37% across two assays. Only donors SCOPE2243 and SCOPE1597 showed an increase of over 0.5 logs, but these increases were not observed consistently across multiple assays. The variability may reflect the fact that the frequency of intact proviruses for these donors was close to the limit of detection based on proviral frequency and the number of cells available for the assay. Due to the low frequency of cells harboring intact provirus, large volumes of blood are required for each assay, limiting the number of repeat experiments for individual participants. Our study compared participants on short-term and long-term ART and found no correlation between resistance to killing and time on ART, indicating that no detectable selection for resistant cells occurs over time in people with suppressed viremia. However, the cohort size might have limited our ability to detect patterns related to treatment duration, time of treatment initiation, different reservoir sizes, and treatment regimens. Increasing cohort size could help identify outliers or patterns and strengthen the conclusions of this study. In addition, the scDbs used in our co-culture assays trigger CTL activation through direct binding to CD3, bypassing the need for TCR engagement in a physiological setting. This means that potential resistance mechanism that rely on the evasion of TCR mechanosensing would not be evident in our assays. Furthermore, the nature of this assay, requiring DNA isolation for the IPDA, prevents us from performing additional single-cell analysis after the co-culture. We are unable to assess differences between CD4^+^ T cell subpopulations such as naïve, central memory and effector memory T cells, which express different levels of Bcl-2, and which were previously reported to exhibit differential susceptibility to killing (91). We also could not perform HIV-1 envelope or integration site sequencing, and therefore cannot assess the composition of each participant’s reservoir and how clonality correlates with observed resistance to killing. The presence of a large clone with increased resistance to CTL-mediated lysis could be a possible explanation for the observed enrichment for cells harboring intact HIV-1 provirus in study participants CW111 and SCOPE2243. Future experiments assessing the composition of the HIV-1 reservoir within study participants and correlating it with the overall or clone-dependent susceptibility to CTL-mediated lysis would expand on our findings and further inform efforts to target the HIV-1 reservoir.

Nonetheless, our study shows that under identical CTL-induced pressure, CD4^+^ T cells from PLWH do not show general increased resistance to cell death regardless of time on ART, indicating that previously reported phenotypic differences between CD4^+^ T cell subpopulations do not significantly limit their susceptibility to cytolytic effector mechanisms. We further show that actively infected cells can partially evade CTL responses restricted to HLA-A2 – and likely also other classical MHC class I molecules – through Nef-dependent reduction of surface antigen availability, but are otherwise equally susceptible to diabody-triggered killing as their uninfected counterparts.

## Materials & Methods

### Sex as a biological variable

Sex was not considered as a biological variable. Both sexes were represented, although most participants were male. Our findings are expected to be relevant to both sexes.

### Study Participants

12 study participants were included in the ex vivo arm of this study. Criteria for the inclusion were expression of HLA-A*02:01 and continuous suppression of HIV-1 viremia through combination antiretroviral treatment for at least 1 year. We received deidentified leukapheresis samples from 9 study participants from the UCSF SCOPE cohort, 2 samples from the University of Toronto and 1 participant from the Bartlett Specialty Clinic at the Johns Hopkins University. Viral loads, CD4^+^ T cell counts and treatment regimens can be found in Supplementary Figure 1.

Healthy donor leukapheresis samples of donors expressing HLA-A*02:01 were purchased from StemCell.

### Cell culture and reagents

PBMCs were isolated using previously a described Ficoll isolation protocol, viably frozen in 90% Fetal Bovine Serum (FBS, GeminiBio, Cat. # 100-800) and 10% DMSO (Sigma-Aldrich, Cat. # 41639) and stored in liquid nitrogen. Cells were thawed in 10mL base media (RPMI1640+Glutamax (FisherScientific, Cat. # 61-870-127) + 10% FBS + 1% Penicillin/Streptomycin (FisherScientific, Cat. # 15140122)) and rested for 3-24 hours before further processing.

Effector cells for co-cultures were expanded through stimulation with soluble anti-CD3 mAb (15 ng/mL, clone OKT3, BioLegend, Cat. # 317302) in CD8 media (base media + 250 U/uL human IL-2 (R&D Systems, Cat. # 202-IL-050/CF) + 5 ng/mL recombinant human IL-7 (BioLegend, Cat. # 581906)) for 72 hours. Cells were then washed and rested in CD8 media for 3 days before CD8^+^ T cell isolation using the CD8^+^ T cell Enrichment Kit (StemCell, Cat. # 19053). Cells were then rested for an additional 4 days before use in co-cultures.

CD4^+^ T cells were cultured in T Cell Growth Media (base media + 100 U/mL recombinant human IL-2 + 1% T Cell Growth Factor (92)), and cells isolated from PLWH were additionally supplemented with Emtricitabine (10 μM FTC, Selleck Chemicals, Cat. # S1704) and Tenofovir (10μM TDF, Sigma-Aldrich, Cat. # 1643656) for the entire duration of culture and co-culture.

The HLA-A2:p53-specific scDb was previously described in (71). The HLA-E:Mtb44-specific scDb was designed and validated in-house (manuscript in preparation). Protein expression and purification was performed by GeneArt (ThermoFisher).

Peptides were synthesized at >95% purity at Elim Biopharm.

### Viral construct and virus production

The NL4.3 ΔEnv-eGFP backbone expression plasmid was obtained through the NIH HIV Reagent Program, Division of AIDS, NIAID, NIH: Human Immunodeficiency Virus 1 (HIV-1) NL4-3 ΔEnv EGFP Reporter Vector, ARP-11100, contributed by Dr. Haili Zhang, Dr. Yan Zhou and Dr. Robert Siliciano. The HIV env expression vector was obtained through the NIH HIV Reagent Program, Division of AIDS, NIAID, NIH: Human Immunodeficiency Virus-1 89.6 Env Expression Vector (pcDNA 89.6 env), ARP-12485, contributed by Dr. Kathleen Collins and Dr. Ronald Collman. The NL4-3 Δnef-eGFP expression vector was constructed as previously described (93). The NL4.3 IRES-eGFP expression plasmid was obtained through the NIH HIV Reagent Program, Division of AIDS, NIAID, NIH:

Human Immunodeficiency Virus Type 1 (HIV-1) NL4-3 IRES-eGFP Infectious Molecular Clone (pBR43IeG-nef+), ARP-11349, contributed by Dr. Jan Münch, Dr. Michael Schindler and Dr. Frank Kirchhoff.

Virus was generated by co-transfecting HEK293T cells in 150cm^2^ cell culture flasks with plasmids encoding for backbone (40μg), HIV envelope (strain 89.6; 20μg; only for envelope-deficient backbone) and supplementary vector pAdvantage (4μg; Promega Cat. # E1711) using the Lipofectamine 3000 Transfection Reagent kit (ThermoScientific, Cat. L3000001) according to manufacturer instructions. Virus-containing supernatant was harvested 48 hours and 72 hours post-transfection, and virus was isolated using a 0.22μm filter and ultracentrifugation with 20% sucrose. Isolated virus was then titered using a p24 ELISA kit (PerkinElmer) following the manufacturer protocol.

### Co-culture killing assays

For ex vivo co-cultures, CD4^+^ T cells from PLWH were isolated using the EasySep Human CD4^+^ T Cell Enrichment Kit (StemCell, Cat. # 19052) according to manufacturer protocol. Cells were then pulsed at a density of 2x10^6^ cells/mL in RPMI media (Gibco, Cat. # 61870036) supplemented with 2% Fetal Bovine Serum and 20μM p53^R175H^ for 4 hours at 37°C and 5% CO_2_. Cells were then washed and resuspended at 1x10^6^ cells/mL in T Cell Growth Media. Autologous CTLs were prepared as described above and resuspended at the same concentration in T Cell Growth Media. CTLs and peptide-pulsed CD4^+^ T cells were plated at effector:target ratios of 1:1-2.5 in UltraLow Attachment 24-well plates (Corning, Cat. # 3473) with 1nM p53-specific scDb. After 18 hours at 37°C, cells were pooled, and viable CD4^+^ T cells were re-isolated through the Milteniy Dead Cell Removal Kit (Cat. # 130-090-101) followed by the EasySep human CD4^+^ T cell Enrichment Kit. Genomic DNA was isolated for further use in the IPDA using the QIAamp DNA Mini Kit (QIAGEN, Cat. # 51304).

For co-cultures using HIV reporter constructs, healthy donor PBMCs were stimulated with 0.5mg/mL PHA in T Cell Growth Media for 72 hours, and then infected through centrifugation with the respective reporter virus for 2 hours at 30°C and 1,200x g. Cells were cultured for an additional 72 hours before peptide pulsing as described above, with 25 µM of either *p53^R175H^* or *Mtb44*. Cells were then plated with autologous CTLs at an effector:target ratio of 1:1 in 96-well V-bottom plates and 20 µL Precision Count Beads per well for 18 hours before analysis via flow cytometry.

### Viral Quantitation Assay

Primary CD4^+^ T cells were isolated from frozen PBMCs from study participants using the StemCell CD4^+^ T cell Enrichment Kit (Cat. # 19052) following the manufacturer protocol, and pulsed with 20 μM of either irrelevant peptide (HLA-A2 signal peptide, VMAPRTLVL) or p53^R175H^ target peptide (HMTEVVRHC) as described above. Cells were plated at a final density of 5x10^6^ cells/mL in T Cell Growth Media in 24-well plates (Corning, Cat. # 3473) at 1 mL per well. p53-specific scDb at a final concentration of 5nM, or PMA/I (50 ng/mL PMA, 1 μM ionomycin) was added to the respective wells, and cells were incubated at 37°C and 5% CO_2_ for 18 hours. Three technical replicates were prepared for each condition. Next, cells were counted and RNA was isolated using the Zymo Quick-RNA viral kit (Cat, # R1034) following the manufacturer protocol, including the Proteinase K and DNase I treatment steps. Extracted RNA was quantified using Qubit measurement (Qubit RNA BR reagent kit, ThermoFisher, Cat. # Q10210) and converted into cDNA using oligo-dT and random hexamer primers using Induro Reverse Transcriptase (NEB, Cat. # M0681S) including RNaseOUT (ThermoFisher, Cat. # 10777019). The mix of primers and RNA was first denatured at 65°C for 5 minutes before adding the reverse transcriptase mix. The reverse transcription reaction occurred with the following thermocycling conditions:

| Temperature (Celsius) | Duration (mm:ss) |
| --- | --- |
| 25 | 5:00 |
| 55 | 30:00 |
| 95 | 1:00 |
| 4 | hold |

For subsequent reactions, cDNA was diluted 2-fold with H_2_O. Three RNA species (poly-adenylated HIV-1 RNA (75, 94), HIV-1 integrase RNA (95), and vpu-envelope spliced HIV-1 RNA (96)) were measured using previously published primers and probes. Probes were double quenched with Zen and Iowa black (Integrated DNA Technologies) and labeled with FAM, Texas Red, and Hex, respectively. Digital PCR reactions were performed on the QIAcuity dPCR machine. Cycling conditions were the following:

| Temperature (Celsius) | Duration (mm:ss) |  |
| --- | --- | --- |
| 95 | 2:00 |  |
| 95 | 0:15 | 45x |
| 56 | 0:30 |  |
| 4 | hold |  |

The RNA from each culture well supernatant was measured as the average of two technical replicates of the dPCR reaction.

### Intact Proviral DNA Assay (IPDA)

To quantify the frequency of intact provirus before and after co-cultures, viable CD4^+^ T cells were isolated using the Milteniy Dead Cell Removal kit (Cat. No# 130-090-101) and the StemCell EasySep Human CD4^+^ T cell Enrichment kit (Cat. No# 19052), and genomic DNA was isolated with the Qiagen QIAamp DNA Micro kit (Cat. No# 56304) and used in the IPDA as previously described (70).

### Flow cytometry

All flow cytometry experiments were performed on a Cytek Northern Lights spectral flow cytometer (3L V-B-R configuration) using the SpectroFlo software (version 3.3.0). Single-color stained controls were included in all experiments and used for the unmixing algorithm included in the software. Cells were stained using the following antibodies purchased from BioLegend at a 70-fold dilution in PBS: APC-conjugated anti-human CD69 (clone FN50, Cat. No# 310910), Brilliant Violet 421-conjugated anti-human CD3 (clone OKT3, Cat. No# 317344), Brilliant Violet 605-conjugated anti-human CD8 (clone SK1, Cat. No# 344742) and, for the experiments in Figure 4A, PE-conjugated anti-human HLA-A2 (Cat. No# 343306) or PE-conjugated anti-human HLA-E (Cat. No# 342604). Additionally, we used eBioscience Fixable Viability Dye eFluor780 (Invitrogen, Cat. No# 65-0865-14) at a 1,000-fold dilution.

### Data analysis

Flow cytometry data was analyzed using FloJo (BD, v10.10.0). For ex vivo co-cultures using cells isolated from PLWH, the frequency of viable CD3^+^CD8^-^ cells was normalized to that of the “no scDb”-control. For co-cultures using healthy donor cells infected with eGFP-reporter virus, the viability of GFP^-^ and GFP^+^ target cells was calculated by normalizing the cell-per-bead ratio for the killing condition to that of the control scDb condition. For example, when targeting an HLA-A2 antigen, the cell-per-bead ratio in the p53-specific scDb-containing well would be normalized to the cell-per-bead ratio in the Mtb44-specific scDb-containing control well.

Digital droplet PCR data was analyzed using QX Manager (BioRad, v 2.1.0).

### Statistical analysis

Statistical significance was calculated with the 2-tailed Student’s t test using GraphPad Prism 10.0. A *P* value of 0.05 was considered significant.

### Study approval

The Institutional Review Boards of UCSF (SCOPE), Johns Hopkins University Bartlett Clinic, and University of Toronto (Unity Health Toronto) approved the study. All participants provided written informed consent.

**Table 1:** Characteristics of study participants. Abbreviations: AA, African American; 3TC, lamivudine; ABC, abacavir; BIC, bictegravir; DOR, doravirine; FTC, emtricitabine; TAF, tenofovir alafenamide; TDF, tenofovir disoproxil fumarate; RPV, rilpivirine; RGV, raltegravir; NVP, nevirapine; TCV, dolutegravir; EGV, elvitegravir; COBI, cobicistat; ATV, atazanavir.

| PID | Age | Sex | Ethnicity | Current ART Regimen | ART initiation | Years on ART | CD4 nadir (cells/μL) | CD4 count (cells/μL) | Plasma HIV-1 RNA (copies/mL) |
| --- | --- | --- | --- | --- | --- | --- | --- | --- | --- |
| SCOPE1151 | 53 | M | Caucasian | BIC/FTC/TAF | Early | 8.9 | 502 | 1017 | <40 |
| SCOPE1597 | 59 | M | Mixed | RPV/TAF/FTC | Chronic | 5.3 | 335 | 350 | <30 |
| SCOPE1713 | 40 | M | Caucasian | BIC/FTC/TAF | Early | 13.2 | 50 | 498 | <40 |
| SCOPE2238 | 67 | M | Caucasian | FTC/TDF, RGV | Early | 9.8 | 538 | 833 | <40 |
| SCOPE2243 | 49 | M | Caucasian | NVP, TDF/FTC | Early | 7.1 | 540 | 806 | <40 |
| SCOPE2303 | 45 | M | Caucasian | ABC/TCV/3TC | Early | 9.7 | 478 | 807 | <40 |
| SCOPE2594 | 30 | M | Hispanic | EGV/TAF/FTC/COBI | Early | 3.6 | 609 | 1405 | <40 |
| SCOPE2647 | 35 | M | Hispanic | EGV/TAF/FTC/COBI | Early | 6.1 | 486 | 654 | <40 |
| SCOPE3054 | 55 | M | Hispanic | FTC, ATV, RGV | Chronic | 5.7 | 9 | 487 | <40 |
| OM5418 | 59 | M | Caucasian | BIG/FTC/TAF, DOR | Chronic | 26.3 | 270 | 616 | <40 |
| CW111 | 67 | F | AA | BIC/FTC/TAF | Chronic | 17.9 | 133 | 874 | <20 |
| CIRC0226 | 56 | M | Hispanic | TCV/3TC | chronic | 22.7 | 8 | 458 | <40 |

## Data availability

Data for the figures are in the Supporting Data Values file.

## Author contributions

N. B. and R. F. S. conceived the study and designed experiments. N. B., B. K. and F. R. S. conducted experiments. H. Z. performed cell sorting. C. K., S. D. and R. H. provided clinical samples and patient data. N. B. and R. F. S. analyzed data and interpreted results. N. B. and R. F. S. wrote the manuscript with input from J. D. S. All authors reviewed, edited and approved the final manuscript.

## Acknowledgements

We want to thank all study participants for contributing to this study. We also want to thank the clinical staff at UCSF, the Maple Leaf Clinic in Toronto and the Bartlett Specialty Clinic at the Johns Hopkins Hospital in Baltimore.

## References

1. Organization WH. HIV and AIDS. https://www.who.int/news-room/fact-sheets/detail/hiv-aids. Updated 15 July 2025 Accessed 15 October, 2025.

2. Blattner W, Gallo RC, and Temin HM. HIV causes AIDS. Science. 1988;241(4865):515–6.

3. Perelson AS, Essunger P, Cao Y, Vesanen M, Hurley A, Saksela K, et al. Decay characteristics of HIV-1-infected compartments during combination therapy. Nature. 1997;387(6629):188–91.

4. Gulick RM, Mellors JW, Havlir D, Eron JJ, Gonzalez C, McMahon D, et al. Treatment with indinavir, zidovudine, and lamivudine in adults with human immunodeficiency virus infection and prior antiretroviral therapy. N Engl J Med. 1997;337(11):734–9.

5. Hammer SM, Squires KE, Hughes MD, Grimes JM, Demeter LM, Currier JS, et al. A controlled trial of two nucleoside analogues plus indinavir in persons with human immunodeficiency virus infection and CD4 cell counts of 200 per cubic millimeter or less. AIDS Clinical Trials Group 320 Study Team. N Engl J Med. 1997;337(11):725–33.

6. Chun TW, Stuyver L, Mizell SB, Ehler LA, Mican JA, Baseler M, et al. Presence of an inducible HIV-1 latent reservoir during highly active antiretroviral therapy. Proc Natl Acad Sci U S A. 1997;94(24):13193–7.

7. Chun TW, Carruth L, Finzi D, Shen X, DiGiuseppe JA, Taylor H, et al. Quantification of latent tissue reservoirs and total body viral load in HIV-1 infection. Nature. 1997;387(6629):183–8.

8. Finzi D, Hermankova M, Pierson T, Carruth LM, Buck C, Chaisson RE, et al. Identification of a reservoir for HIV-1 in patients on highly active antiretroviral therapy. Science. 1997;278(5341):1295–300.

9. Wong JK, Hezareh M, Günthard HF, Havlir DV, Ignacio CC, Spina CA, et al. Recovery of replication-competent HIV despite prolonged suppression of plasma viremia. Science. 1997;278(5341):1291–5.

10. Siliciano JD, Kajdas J, Finzi D, Quinn TC, Chadwick K, Margolick JB, et al. Long-term follow-up studies confirm the stability of the latent reservoir for HIV-1 in resting CD4+ T cells. Nat Med. 2003;9(6):727–8.

11. Crooks AM, Bateson R, Cope AB, Dahl NP, Griggs MK, Kuruc JD, et al. Precise Quantitation of the Latent HIV-1 Reservoir: Implications for Eradication Strategies. J Infect Dis. 2015;212(9):1361–5.

12. McMyn NF, Varriale J, Fray EJ, Zitzmann C, MacLeod H, Lai J, et al. The latent reservoir of inducible, infectious HIV-1 does not decrease despite decades of antiretroviral therapy. J Clin Invest. 2023;133(17).

13. Strain MC, Günthard HF, Havlir DV, Ignacio CC, Smith DM, Leigh-Brown AJ, et al. Heterogeneous clearance rates of long-lived lymphocytes infected with HIV: intrinsic stability predicts lifelong persistence. Proc Natl Acad Sci U S A. 2003;100(8):4819–24.

14. Chun TW, Davey RT, Engel D, Lane HC, and Fauci AS. Re-emergence of HIV after stopping therapy. Nature. 1999;401(6756):874–5.

15. Rothenberger MK, Keele BF, Wietgrefe SW, Fletcher CV, Beilman GJ, Chipman JG, et al. Large number of rebounding/founder HIV variants emerge from multifocal infection in lymphatic tissues after treatment interruption. Proc Natl Acad Sci U S A. 2015;112(10):E1126–34.

16. Gunst JD, Gohil J, Li JZ, Bosch RJ, White Catherine Seamon A, Chun TW, et al. Time to HIV viral rebound and frequency of post-treatment control after analytical interruption of antiretroviral therapy: an individual data-based meta-analysis of 24 prospective studies. Nat Commun. 2025;16(1):906.

17. Collins DR, Gaiha GD, and Walker BD. CD8. Nat Rev Immunol. 2020;20(8):471–82.

18. Walker BD, Chakrabarti S, Moss B, Paradis TJ, Flynn T, Durno AG, et al. HIV-specific cytotoxic T lymphocytes in seropositive individuals. Nature. 1987;328(6128):345–8.

19. Kwaa AK, and Blankson JN. Immune Responses in Controllers of HIV Infection. Annu Rev Immunol. 2024;42(1):21–33.

20. Koup RA, Safrit JT, Cao Y, Andrews CA, McLeod G, Borkowsky W, et al. Temporal association of cellular immune responses with the initial control of viremia in primary human immunodeficiency virus type 1 syndrome. J Virol. 1994;68(7):4650–5.

21. Borrow P, Lewicki H, Hahn BH, Shaw GM, and Oldstone MB. Virus-specific CD8+ cytotoxic T-lymphocyte activity associated with control of viremia in primary human immunodeficiency virus type 1 infection. J Virol. 1994;68(9):6103–10.

22. Schmitz JE, Kuroda MJ, Santra S, Sasseville VG, Simon MA, Lifton MA, et al. Control of viremia in simian immunodeficiency virus infection by CD8+ lymphocytes. Science. 1999;283(5403):857–60.

23. Jin X, Bauer DE, Tuttleton SE, Lewin S, Gettie A, Blanchard J, et al. Dramatic rise in plasma viremia after CD8(+) T cell depletion in simian immunodeficiency virus-infected macaques. J Exp Med. 1999;189(6):991–8.

24. Migueles SA, Sabbaghian MS, Shupert WL, Bettinotti MP, Marincola FM, Martino L, et al. HLA B*5701 is highly associated with restriction of virus replication in a subgroup of HIV-infected long term nonprogressors. Proc Natl Acad Sci U S A. 2000;97(6):2709–14.

25. Pereyra F, Jia X, McLaren PJ, Telenti A, de Bakker PI, Walker BD, et al. The major genetic determinants of HIV-1 control affect HLA class I peptide presentation. Science. 2010;330(6010):1551–7.

26. Archin NM, Liberty AL, Kashuba AD, Choudhary SK, Kuruc JD, Crooks AM, et al. Administration of vorinostat disrupts HIV-1 latency in patients on antiretroviral therapy. Nature. 2012;487(7408):482–5.

27. Deeks SG. HIV: Shock and kill. Nature. 2012;487(7408):439–40.

28. Westendorp MO, Frank R, Ochsenbauer C, Stricker K, Dhein J, Walczak H, et al. Sensitization of T cells to CD95-mediated apoptosis by HIV-1 Tat and gp120. Nature. 1995;375(6531):497–500.

29. Bartz SR, and Emerman M. Human immunodeficiency virus type 1 Tat induces apoptosis and increases sensitivity to apoptotic signals by up-regulating FLICE/caspase-8. J Virol. 1999;73(3):1956–63.

30. Thakur BK, Chandra A, Dittrich T, Welte K, and Chandra P. Inhibition of SIRT1 by HIV-1 viral protein Tat results in activation of p53 pathway. Biochem Biophys Res Commun. 2012;424(2):245–50.

31. Dabrowska A, Kim N, and Aldovini A. Tat-induced FOXO3a is a key mediator of apoptosis in HIV-1-infected human CD4+ T lymphocytes. J Immunol. 2008;181(12):8460–77.

32. López-Huertas MR, Mateos E, Sánchez Del Cojo M, Gómez-Esquer F, Díaz-Gil G, Rodríguez-Mora S, et al. The presence of HIV-1 Tat protein second exon delays fas protein-mediated apoptosis in CD4+ T lymphocytes: a potential mechanism for persistent viral production. J Biol Chem. 2013;288(11):7626–44.

33. Zauli G, Gibellini D, Caputo A, Bassini A, Negrini M, Monne M, et al. The human immunodeficiency virus type-1 Tat protein upregulates Bcl-2 gene expression in Jurkat T-cell lines and primary peripheral blood mononuclear cells. Blood. 1995;86(10):3823–34.

34. Zauli G, and Gibellini D. The human immunodeficiency virus type-1 (HIV-1) Tat protein and Bcl-2 gene expression. Leuk Lymphoma. 1996;23(5-6):551–60.

35. Chandrasekar AP, Cummins NW, and Badley AD. The Role of the BCL-2 Family of Proteins in HIV-1 Pathogenesis and Persistence. Clin Microbiol Rev. 2019;33(1).

36. Geleziunas R, Xu W, Takeda K, Ichijo H, and Greene WC. HIV-1 Nef inhibits ASK1-dependent death signalling providing a potential mechanism for protecting the infected host cell. Nature. 2001;410(6830):834–8.

37. Kumar B, Tripathi C, Kanchan RK, Tripathi JK, Ghosh JK, Ramachandran R, et al. Dynamics of physical interaction between HIV-1 Nef and ASK1: identifying the interacting motif(s). PLoS One. 2013;8(6):e67586.

38. Sevilya Z, Chorin E, Gal-Garber O, Zelinger E, Turner D, Avidor B, et al. Killing of Latently HIV-Infected CD4 T Cells by Autologous CD8 T Cells Is Modulated by Nef. Front Immunol. 2018;9:2068.

39. Wolf D, Witte V, Laffert B, Blume K, Stromer E, Trapp S, et al. HIV-1 Nef associated PAK and PI3-kinases stimulate Akt-independent Bad-phosphorylation to induce anti-apoptotic signals. Nat Med. 2001;7(11):1217–24.

40. Greenway AL, McPhee DA, Allen K, Johnstone R, Holloway G, Mills J, et al. Human immunodeficiency virus type 1 Nef binds to tumor suppressor p53 and protects cells against p53-mediated apoptosis. J Virol. 2002;76(6):2692–702.

41. Percario Z, Olivetta E, Fiorucci G, Mangino G, Peretti S, Romeo G, et al. Human immunodeficiency virus type 1 (HIV-1) Nef activates STAT3 in primary human monocyte/macrophages through the release of soluble factors: involvement of Nef domains interacting with the cell endocytotic machinery. J Leukoc Biol. 2003;74(5):821–32.

42. Ndolo T, Dhillon NK, Nguyen H, Guadalupe M, Mudryj M, and Dandekar S. Simian immunodeficiency virus Nef protein delays the progression of CD4+ T cells through G1/S phase of the cell cycle. J Virol. 2002;76(8):3587–95.

43. Rasola A, Gramaglia D, Boccaccio C, and Comoglio PM. Apoptosis enhancement by the HIV-1 Nef protein. J Immunol. 2001;166(1):81–8.

44. Laforge M, Petit F, Estaquier J, and Senik A. Commitment to apoptosis in CD4(+) T lymphocytes productively infected with human immunodeficiency virus type 1 is initiated by lysosomal membrane permeabilization, itself induced by the isolated expression of the viral protein Nef. J Virol. 2007;81(20):11426–40.

45. Xu XN, Screaton GR, Gotch FM, Dong T, Tan R, Almond N, et al. Evasion of cytotoxic T lymphocyte (CTL) responses by nef-dependent induction of Fas ligand (CD95L) expression on simian immunodeficiency virus-infected cells. J Exp Med. 1997;186(1):7–16.

46. Nie Z, Phenix BN, Lum JJ, Alam A, Lynch DH, Beckett B, et al. HIV-1 protease processes procaspase 8 to cause mitochondrial release of cytochrome c, caspase cleavage and nuclear fragmentation. Cell Death Differ. 2002;9(11):1172–84.

47. Nie Z, Bren GD, Rizza SA, and Badley AD. HIV Protease Cleavage of Procaspase 8 is Necessary for Death of HIV-Infected Cells. Open Virol J. 2008;2:1–7.

48. Sainski AM, Natesampillai S, Cummins NW, Bren GD, Taylor J, Saenz DT, et al. The HIV-1-specific protein Casp8p41 induces death of infected cells through Bax/Bak. J Virol. 2011;85(16):7965–75.

49. Sainski AM, Dai H, Natesampillai S, Pang YP, Bren GD, Cummins NW, et al. Casp8p41 generated by HIV protease kills CD4 T cells through direct Bak activation. J Cell Biol. 2014;206(7):867–76.

50. Nie Z, Bren GD, Vlahakis SR, Schimnich AA, Brenchley JM, Trushin SA, et al. Human immunodeficiency virus type 1 protease cleaves procaspase 8 in vivo. J Virol. 2007;81(13):6947–56.

51. Jacotot E, Ferri KF, El Hamel C, Brenner C, Druillennec S, Hoebeke J, et al. Control of mitochondrial membrane permeabilization by adenine nucleotide translocator interacting with HIV-1 viral protein rR and Bcl-2. J Exp Med. 2001;193(4):509–19.

52. Stewart SA, Poon B, Jowett JB, and Chen IS. Human immunodeficiency virus type 1 Vpr induces apoptosis following cell cycle arrest. J Virol. 1997;71(7):5579–92.

53. Casella CR, Rapaport EL, and Finkel TH. Vpu increases susceptibility of human immunodeficiency virus type 1-infected cells to fas killing. J Virol. 1999;73(1):92–100.

54. Akari H, Bour S, Kao S, Adachi A, and Strebel K. The human immunodeficiency virus type 1 accessory protein Vpu induces apoptosis by suppressing the nuclear factor kappaB-dependent expression of antiapoptotic factors. J Exp Med. 2001;194(9):1299–311.

55. Airò P, Torti C, Uccelli MC, Malacarne F, Palvarini L, Carosi G, et al. I CD4+ T lymphocytes express high levels of Bcl-2 after highly active antiretroviral therapy for HIV infection. AIDS Res Hum Retroviruses. 2000;16(17):1805–7.

56. Ren Y, Huang SH, Patel S, Alberto WDC, Magat D, Ahimovic D, et al. BCL-2 antagonism sensitizes cytotoxic T cell-resistant HIV reservoirs to elimination ex vivo. J Clin Invest. 2020;130(5):2542–59.

57. McGary CS, Deleage C, Harper J, Micci L, Ribeiro SP, Paganini S, et al. CTLA-4. Immunity. 2017;47(4):776–88.e5.

58. Gattinoni L, Lugli E, Ji Y, Pos Z, Paulos CM, Quigley MF, et al. A human memory T cell subset with stem cell-like properties. Nat Med. 2011;17(10):1290–7.

59. Buzon MJ, Sun H, Li C, Shaw A, Seiss K, Ouyang Z, et al. HIV-1 persistence in CD4+ T cells with stem cell-like properties. Nat Med. 2014;20(2):139–42.

60. Palmer CS, Duette GA, Wagner MCE, Henstridge DC, Saleh S, Pereira C, et al. Metabolically active CD4+ T cells expressing Glut1 and OX40 preferentially harbor HIV during in vitro infection. FEBS Lett. 2017;591(20):3319–32.

61. Huang SH, Ren Y, Thomas AS, Chan D, Mueller S, Ward AR, et al. Latent HIV reservoirs exhibit inherent resistance to elimination by CD8+ T cells. J Clin Invest. 2018;128(2):876–89.

62. Bruner KM, Murray AJ, Pollack RA, Soliman MG, Laskey SB, Capoferri AA, et al. Defective proviruses rapidly accumulate during acute HIV-1 infection. Nat Med. 2016;22(9):1043–9.

63. Pollack RA, Jones RB, Pertea M, Bruner KM, Martin AR, Thomas AS, et al. Defective HIV-1 Proviruses Are Expressed and Can Be Recognized by Cytotoxic T Lymphocytes, which Shape the Proviral Landscape. Cell Host Microbe. 2017;21(4):494–506.e4.

64. Ferreira IATM, Herrera A, Huynh TT, Stone E, Linden NL, Ovies C, et al. Dynamic antigen expression and cytotoxic T cell resistance in HIV reservoir clones. Nature. 2026.

65. Leyre L, Mustapha F, Herrera A, Lee E, Huntsman E, Zumbo P, et al. HIV Nef amplifies mechanical heterogeneity to promote immune evasion. bioRxiv. 2025.

66. Lian X, Seiger KW, Parsons EM, Gao C, Sun W, Gladkov GT, et al. Progressive transformation of the HIV-1 reservoir cell profile over two decades of antiviral therapy. Cell Host Microbe. 2023;31(1):83–96.e5.

67. McMyn NF, Varriale J, Wu HWS, Hariharan V, Moskovljevic M, Tan TS, et al. Factors associated with resistance of HIV-1 reservoir viruses to neutralization by autologous IgG antibodies. J Clin Invest. 2025;135(19).

68. Chun TW, Finzi D, Margolick J, Chadwick K, Schwartz D, and Siliciano RF. In vivo fate of HIV-1-infected T cells: quantitative analysis of the transition to stable latency. Nat Med. 1995;1(12):1284–90.

69. Shan L, Deng K, Gao H, Xing S, Capoferri AA, Durand CM, et al. Transcriptional Reprogramming during Effector-to-Memory Transition Renders CD4. Immunity. 2017;47(4):766–75.e3.

70. Bruner KM, Wang Z, Simonetti FR, Bender AM, Kwon KJ, Sengupta S, et al. A quantitative approach for measuring the reservoir of latent HIV-1 proviruses. Nature. 2019;566(7742):120–5.

71. Hsiue EH, Wright KM, Douglass J, Hwang MS, Mog BJ, Pearlman AH, et al. Targeting a neoantigen derived from a common. Science. 2021;371(6533).

72. Simonetti FR, White JA, Tumiotto C, Ritter KD, Cai M, Gandhi RT, et al. Intact proviral DNA assay analysis of large cohorts of people with HIV provides a benchmark for the frequency and composition of persistent proviral DNA. Proc Natl Acad Sci U S A. 2020;117(31):18692–700.

73. Siliciano JD, and Siliciano RF. In Vivo Dynamics of the Latent Reservoir for HIV-1: New Insights and Implications for Cure. Annu Rev Pathol. 2022;17:271–94.

74. Moskovljevic M, Dragoni F, Board NL, Wu F, Lai J, Zhang H, et al. Cognate antigen engagement induces HIV-1 expression in latently infected CD4. Immunity. 2024;57(12):2928–44.e6.

75. Shan L, Rabi SA, Laird GM, Eisele EE, Zhang H, Margolick JB, et al. A novel PCR assay for quantification of HIV-1 RNA. J Virol. 2013;87(11):6521–5.

76. Eriksson S, Graf EH, Dahl V, Strain MC, Yukl SA, Lysenko ES, et al. Comparative analysis of measures of viral reservoirs in HIV-1 eradication studies. PLoS Pathog. 2013;9(2):e1003174.

77. Timmons A, Fray E, Kumar M, Wu F, Dai W, Bullen CK, et al. HSF1 inhibition attenuates HIV-1 latency reversal mediated by several candidate LRAs In Vitro and Ex Vivo. Proc Natl Acad Sci U S A. 2020;117(27):15763–71.

78. Hosmane NN, Kwon KJ, Bruner KM, Capoferri AA, Beg S, Rosenbloom DI, et al. Proliferation of latently infected CD4. J Exp Med. 2017;214(4):959–72.

79. Kwon KJ, Timmons AE, Sengupta S, Simonetti FR, Zhang H, Hoh R, et al. Different human resting memory CD4. Sci Transl Med. 2020;12(528).

80. Ho YC, Shan L, Hosmane NN, Wang J, Laskey SB, Rosenbloom DI, et al. Replication-competent noninduced proviruses in the latent reservoir increase barrier to HIV-1 cure. Cell. 2013;155(3):540–51.

81. Collins KL, Chen BK, Kalams SA, Walker BD, and Baltimore D. HIV-1 Nef protein protects infected primary cells against killing by cytotoxic T lymphocytes. Nature. 1998;391(6665):397–401.

82. Blagoveshchenskaya AD, Thomas L, Feliciangeli SF, Hung CH, and Thomas G. HIV-1 Nef downregulates MHC-I by a PACS-1- and PI3K-regulated ARF6 endocytic pathway. Cell. 2002;111(6):853–66.

83. Collins KL. Resistance of HIV-infected cells to cytotoxic T lymphocytes. Microbes Infect. 2004;6(5):494–500.

84. Perica K, Kotchetkov IS, Mansilla-Soto J, Ehrich F, Herrera K, Shi Y, et al. HIV immune evasin Nef enhances allogeneic CAR T cell potency. Nature. 2025;640(8059):793–801.

85. Walters LC, Harlos K, Brackenridge S, Rozbesky D, Barrett JR, Jain V, et al. Pathogen-derived HLA-E bound epitopes reveal broad primary anchor pocket tolerability and conformationally malleable peptide binding. Nat Commun. 2018;9(1):3137.

86. Hersperger AR, Migueles SA, Betts MR, and Connors M. Qualitative features of the HIV-specific CD8+ T-cell response associated with immunologic control. Curr Opin HIV AIDS. 2011;6(3):169–73.

87. Dobmeyer TS, Klein SA, Dobmeyer JM, Raffel B, Findhammer S, Hoelzer D, et al. Differential expression of bcl-2 and susceptibility to programmed cell death in lymphocytes of HIV-1-infected individuals. Clin Immunol Immunopathol. 1998;87(3):230–9.

88. Chandrasekar AP, Cummins NW, Natesampillai S, Misra A, Alto A, Laird G, et al. The BCL-2 Inhibitor Venetoclax Augments Immune Effector Function Mediated by Fas Ligand, TRAIL, and Perforin/Granzyme B, Resulting in Reduced Plasma Viremia and Decreased HIV Reservoir Size during Acute HIV Infection in a Humanized Mouse Model. J Virol. 2022;96(24):e0173022.

89. Brazin KN, Mallis RJ, Boeszoermenyi A, Feng Y, Yoshizawa A, Reche PA, et al. The T Cell Antigen Receptor α Transmembrane Domain Coordinates Triggering through Regulation of Bilayer Immersion and CD3 Subunit Associations. Immunity. 2018;49(5):829–41.e6.

90. Kim ST, Takeuchi K, Sun ZY, Touma M, Castro CE, Fahmy A, et al. The alphabeta T cell receptor is an anisotropic mechanosensor. J Biol Chem. 2009;284(45):31028–37.

91. Liu J, and Roederer M. Differential susceptibility of leukocyte subsets to cytotoxic T cell killing: implications for HIV immunopathogenesis. Cytometry A. 2007;71(2):94–104.

92. Siliciano JD, and Siliciano RF. Enhanced culture assay for detection and quantitation of latently infected, resting CD4+ T-cells carrying replication-competent virus in HIV-1-infected individuals. Methods Mol Biol. 2005;304:3–15.

93. O’Connell KA, Rabi SA, Siliciano RF, and Blankson JN. CD4+ T cells from elite suppressors are more susceptible to HIV-1 but produce fewer virions than cells from chronic progressors. Proc Natl Acad Sci U S A. 2011;108(37):E689–98.

94. Bullen CK, Laird GM, Durand CM, Siliciano JD, and Siliciano RF. New ex vivo approaches distinguish effective and ineffective single agents for reversing HIV-1 latency in vivo. Nat Med. 2014;20(4):425–9.

95. Cillo AR, Vagratian D, Bedison MA, Anderson EM, Kearney MF, Fyne E, et al. Improved single-copy assays for quantification of persistent HIV-1 viremia in patients on suppressive antiretroviral therapy. J Clin Microbiol. 2014;52(11):3944–51.

96. Gao H, Ozantürk AN, Wang Q, Harlan GH, Schmitz AJ, Presti RM, et al. Evaluation of HIV-1 latency reversal and antibody-dependent viral clearance by quantification of singly spliced HIV-1. J Virol. 2021;95(11).

