## Supplementary figures and images for "HIV-1 infection does not confer intrinsic resistance to cell death induced by cytotoxic T lymphocytes"

### Supplementary Figure 1

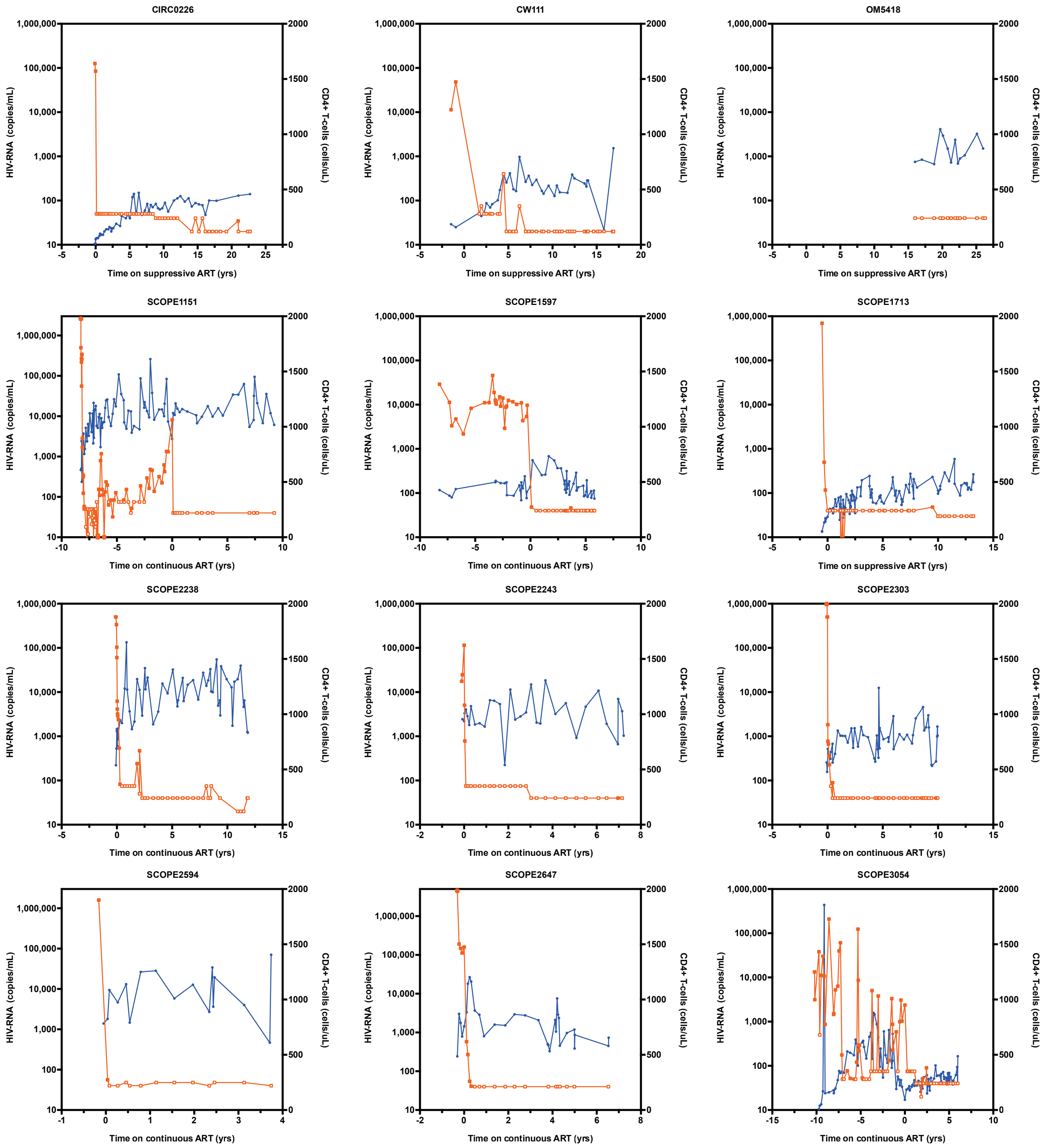
